# Suzetrigine (VX-548) exhibits activity-dependent effects on human dorsal root ganglion neurons

**DOI:** 10.1101/2025.04.09.648014

**Authors:** Megan L. Uhelski, Mandee K. Schaub, Felipe Espinosa, Mario Heles, Nicolas Cortes, Yan Li, Claudio E. Tatsui, Lawrence D. Rhines, Robert Y. North, Christopher Alvarez-Breckendridge, Geoffrey Funk, Peter Horton, Anna Cervantes, Joseph B. Lesnak, Michele Curalto, Gregory Dussor, Theodore J. Price, Patrick M. Dougherty

**Affiliations:** Department of Pain Medicine, University of Texas MD Anderson Cancer Center, Houston, TX, 77030, USA; Department of Neuroscience and Center for Advanced Pain Studies, University of Texas at Dallas, Richardson, TX, 75080, USA; Department of Neurosurgery, University of Texas MD Anderson Cancer Center, Houston, TX, 77030, USA; Southwest Transplant Alliance, Dallas, TX, 75231, USA; Department of Anesthesiology and Pain Medicine, University of Washington, Seattle, WA, 98195, USA

**Keywords:** neuropathy, spontaneous activity, sensitization

## Abstract

Non-selective antagonists of voltage-gated sodium channels (VGSCs) provide effective local analgesia, but systemic administration is fraught with unwanted cardiovascular and central nervous system toxicity. The development of antagonists targeting VGSCs preferentially expressed in peripheral neurons, such as Na_V_1.7, 1.8, and 1.9, could mitigate these risks. One such new small molecule that selectively inhibits Na_V_1.8, is orally bioavailable, and has recent FDA approval for the treatment of acute pain is suzetrigine (VX-548). Here we tested the effects of this agent on human dorsal root ganglion (hDRG) neurons obtained from either patients undergoing surgical thoracic vertebrectomy or from organ donors in parallel electrophysiology studies across 2 laboratories. Bath application of 10 nM suzetrigine abolished spontaneous discharges previously shown to be associated with on-going neuropathic pain within minutes. In contrast, suppression of discharges evoked by intracellular current injection required more prolonged application times. These results suggest that suzetrigine will likely show efficacy versus spontaneous pain but may have less robust effects on evoked or breakthrough pain.

## Introduction

The voltage-gated sodium channel (Na_V_) is an important target for the treatment of pathophysiological conditions associated with aberrant neuronal firing patterns, including pain and epilepsy^1,2^. Unfortunately, non-specific Na_V_ antagonists typically have problematic side effect profiles when used systemically^3,4^. Thus, much interest and effort in pain research has been expended in developing systemically active antagonists specific to the Na_V_ 1.7, 1.8 and 1.9 subtypes preferentially expressed in primary sensory neurons and with well-established roles in pain physiology^5–7^. A breakthrough in this regard is suzetrigine (VX-458) that is specific for the human Na_V_1.8 channel and that was approved in January 2025 by the FDA for acute pain treatment^8–11^. Na_V_1.8 contributes to both neuropathic and inflammatory pain states^12^. Knockdown of Na_V_1.8 in rats reduced tetrodotoxin-resistant (TTX-R) sodium currents and reversed tactile allodynia and hyperalgesia after spinal nerve ligation^13^, while knockout of Na_V_1.8 in mice prevented the development of NGF-induced thermal hyperalgesia^14^. Na_V_1.8 expression increased in non-injured DRG neurons following nerve injury, enhancing TTX-R sodium currents resulting in hyperexcitability and increased AP firing^15^. There is also evidence for delayed increases in Na_V_1.8 expression in injured neurons^16^. Increased Na_V_1.8 expression and enhanced TTX-R sodium currents in rat DRG neurons occurs in response to multiple pro-inflammatory compounds, including PGE_2_, TNF-α, carrageenan, adenosine, and serotonin^17–19^. Finally, gain-of-function mutations in the Na_V_1.8 gene (*SCN10A*) are associated with neuropathic pain conditions^20^.

Studies of Na_V_1.8 blockade in rodent nociceptors are likely to underestimate its effects on human nociceptors in that human Na_V_1.8 shows many key differences in the function including generating larger and more enduring currents; more hyperpolarized activation and slower inactivation relative to membrane potential; faster inactivation at a more depolarized membrane potential; higher potential for spontaneous channel^6^. Indeed, rats expressing human *SCN10A* were much more responsive to a human Nav1.8-selective small molecule inhibitor (MSD199) than wild-type rats in behavioral and neurophysiological assays^21^. Thus, examinations of novel Na_V_1.8 inhibitors specifically designed to target human Na_V_1.8 channels necessitate the use of human or humanized sensory neurons for testing and validation for use in different pain states.

In this work we test the impact of suzetrigine Na_V_1.8 inhibition on human sensory neurons cultured from DRGs both with and without prior pain history^22^. Particular attention was paid to testing effects of Na_V_1.8 inhibition on spontaneous action potential discharges in DRG neurons we previously linked to ongoing neuropathic pain symptoms^22,23^. As well, this work compared the potential impact of tissue source and dissociation methods from both cancer patients with on-going neuropathic pain at the University of Texas MD Anderson Cancer Center (UTMDACC) and from organ donors recovered in collaboration with the Southwest Transplant Alliance (STA) at the University of Texas at Dallas (UTD).

## Materials and Methods

### Subjects

Human tissue procurement procedures were approved by the Institutional Review Boards at UTMDACC and UTD and all experiments conformed to relevant guidelines and regulations and conducted with appropriate written informed consent for participation. Tissue samples from a total of 5 consecutive patients (3 male, 2 female) and 9 organ donors (7 males and 2 females) were used in the study. UTMDACC patient characteristics are presented in Supplementary Table 1. All patient samples were recovered during thoracic vertebrectomy surgeries for relief of physician-documented spinal cord/dorsal root compression. Dermatomes affected by pain were present in all but one patient as defined using pain diagrams collected at the time of consent as in prior studies^22^. STA donor characteristics are presented in Supplementary Tables 2-3 that were obtained from their attendant medical history. Only one donor had prior history of diabetic neuropathy.

### Tissue Preparation

#### MDA Thoracic Vertebrectomy Patients

DRG neurons obtained from patients undergoing surgical treatment that necessitated ligation of spinal nerve roots to facilitate tumor resection and/or spinal reconstruction were dissociated as described previously^22^. Excised DRGs were transferred immediately into cold (∼4°C) and sterile balanced salt solution (EBSS, Gibco) and transported to the laboratory on ice in sterile 50 mL centrifuge tubes. Each ganglion was carefully dissected and divided into ∼1 mm^2^ sections which were moved to a Petri dish containing a 2mL solution of trypsin, collagenase, and DNase (Sigma) in DMEM/F12 (Gibco) and placed in an incubated orbital shaker at 37°C for 20 min. After each 20 min session, the digestion solution was collected, placed into a blocking solution consisting of DMEM/F12 with 10% horse serum (Gibco) and 1% penicillin-streptomycin (Gibco), and replaced with fresh digestion solution. This process was repeated until the sections were adequately digested. The collected digestion solution and blocking solution was centrifuged at 23°C and 180g for 5 min. The supernatant was removed, and the cell pellet re-suspended with fresh culture media consisting of DMEM/F12 with 10% horse serum, 1% penicillin-streptomycin, and 0.1 µg/mL β-NGF (Cell Signaling Technology). Cells were plated on poly-L-lysine-coated (Sigma) round glass coverslips (Chemglass). Detailed recording whole-cell recording protocols are available in Supplemental Materials.

#### UTD STA Donors

DRGs recovered from human organ donors using an established protocol ^24^ were cleaned, dissociated, and cultured for subsequent experiments. Tissue was dissected in aCSF to remove connective tissue and fat from around the bulb. The tissue was then minced and dissociated in 1 mg/mL STEMxyme (Worthington, LS004107) in HBSS. Enzymatic dissociation was done using one of 2 protocols. Protocol 1 was Standard Dissociation, with enzyme solution for 3-5 hours at 37°C, triturating with decreasing sizes of fire polished Pasteur pipettes every hour. Once the tissue was dissociated, the cell solution was run though a 100-µm mesh strainer and then through a 10% BSA gradient centrifuged at 500g for 5 min to remove debris. Cells were then plated on PDL-coated glass coverslips at 50-100 cells per coverslip and used for patch clamp electrophysiology starting after day 3 in vitro. Protocol 2 was dissociation at room temperature (RT). Before mincing, DRGs were weighed, and 1 mL of dissociation solution was used for 15-25 mg of cleaned DRG. The tissue suspension was placed in a 15 mL Falcon tube and constantly mixed on a nutator shaker at 62 rpm. The tissue suspension was monitored closely and, once the tissue became loose (usually within 90-120 min), it was subjected to the trituration/centrifugation steps as described above. The cell pellet was resuspended in culture media. Cells were counted and survival rates were determined with Trypan Blue. Fresh dissociation solution was added to the remaining tissue and subjected to further enzymatic digestion for 30-60 minutes, and the following steps were done as above. Dissociated cells were stored in Eppendorf tube at RT for 0 to 20 days until they were plated. Cells were plated inside cell-culture inserts (IBIDI, Cat# 80209) attached to a 12 mm coverslip (2-4 coverslips per day). Detailed recording whole-cell recording protocols are available in Supplemental Materials.

## Drugs

Suzetrigine (VX-548; MedChemExpress) was prepared in a stock solution of fresh DMSO and diluted in extracellular solution to a working concentration of 10 nM. This concentration was selected based on published studies showing an IC_50_ in the nM range *in vitro*^8,10^.

## Data availability

The data that support the findings of this study are available from the corresponding authors, upon reasonable request.

## Results

A total of 14 neurons from 5 patients and 35 neurons from 9 organ donors (20 from Protocol 1 and 15 from Protocol 2) were tested in the study. Overall, 42% (6) of the patient cells demonstrated spontaneous activity (SA) and all but one of these were from pain-associated dermatomes. Details are provided in Supplementary Table 4. The overall SA incidence was 14% (5) from the organ donor neuron pool but only 1 of these neurons came from a donor with a documented history of neuropathic pain.

### Suzetrigine blocks ectopic action potential discharge in hyperexcitable DRG neurons

The first set of experiments focused on testing whether suzetrigine (VX-548, 10 nM) could suppress on-going action potential (AP) firing. We observed that suzetrigine was quite effective and suppressed AP firing over an interval that varied between 3 to 20 min from the start of drug infusion (Fig.1A-C). This occurred in 6/6 DRG neurons from MDA patients and was also associated with membrane hyperpolarization (Fig.1D). While rheobase was not significantly altered post-drug, AP overshoot (in mV, measuring from the part of the AP waveform that crosses 0 mV to the peak AP amplitude) and AP width at 0 mV decreased post-drug (Fig. 1E-H). No other AP characteristics were significantly altered (Supplementary Fig.1). In contrast, responses to 1s current stimulation at and above rheobase did not differ before and after treatment (Fig.1I). Neurons from STA donor DRGs demonstrating SA at baseline (*n* = 5) were similarly treated with suzetrigine (10 nM) and all showed suppression in this activity over an interval between 3 to 8 min (Fig.1J,K). Suzetrigine decreased RMP in 4 neurons, but one showed marked depolarization as summarized in Fig.1L.

**Figure 1.**
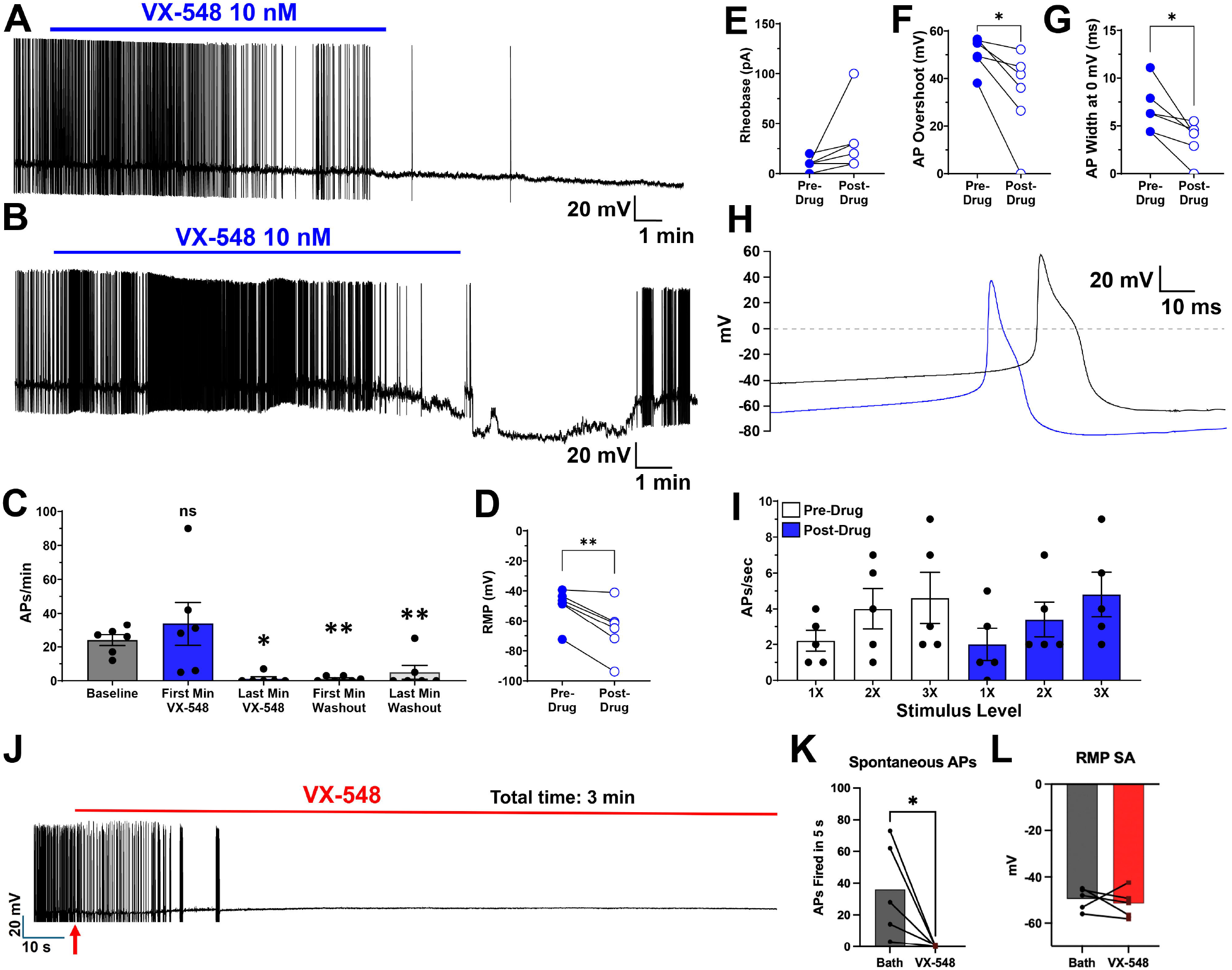
Suzetrigine reversibly inhibits spontaneous action potential discharge in hyperexcitable human DRG neurons from MDA patients and STA donors (Protocol 1). (A) Representative raw trace example showing an MDA patient DRG neuron with ectopic discharge at baseline that ceased action potential (AP) firing and became hyperpolarized after application of 10 nM suzetrigine (VX-548), as indicated by the blue bar above the trace. (B) A second MDA patient DRG neuron raw trace example shows the return of action potential firing after a brief (∼5 min) washout following 10 nM VX-548, as shown by the blue bar above the trace. Summary data for the frequency of AP firing is shown in (C). The firing rate in APs/min increased slightly during the first minute of drug infusion but decreased significantly by the last minute of infusion and remained low during the first minute of washout, *F*(4,20) = 7.71, *P* =.001. At the last minute of washout, AP firing had resumed in two neurons, but at a lower rate than baseline. (D) Suzetrigine led to hyperpolarization of the resting membrane potential (RMP) in all 6 MDA patient DRG neurons, *t*(5) = 5.12, *P* =.004, but as seen in (E), post-drug increases in rheobase were not significant *t*(5) = 1.72, *P* =0.145). An examination of AP characteristics found that (F) AP overshoot *t*(5) = 3.23, *P* =0.023, and (G) AP width at 0 mV, *t*(5) = 3.38, *P* =0.020, were significantly lower post-drug in MDA patient DRG neurons, as demonstrated by the raw trace example in (H), where the pre-drug waveform is shown in black and the post-drug AP waveform is shown in blue. (I) Responses to current stimulation at 1X-3X rheobase did not differ pre- and post-drug in MDA patient DRG neurons, *F*(1,4) = 0.05, *P* =0.831). In DRGs from STA donors, suzetrigine (VX-548) application rapidly abolished AP firing in spontaneously active neurons (*n* = 5), as shown by the raw trace example in (J). Summary data for the frequency of AP firing is shown in (K) and was analyzed using the Wilcoxon matched pairs signed rank test, *P* = 0.0312. As seen in (L), RMP did not change significantly after VX-548 application. \**P*<.05, ** *P*<.01.

### Non-SA neurons from MDA cancer patients exhibited minimal effects following 10 min suzetrigine treatment

Based on the mean time for suppression of on-going SA, neurons from MDA patients without ectopic activity at baseline (non-SA, *n* = 8) were also treated with 10 nM VX-548 for 10 min (Fig. 2A). With this limited treatment time no change in RMP, rheobase, or responses to current stimulation at and above rheobase were found (Fig.2B-D). Suzetrigine did decrease AP overshoot and amplitude (Fig.2E-G), but no other AP characteristics were altered (Supplementary Fig.2).

**Figure 2.**
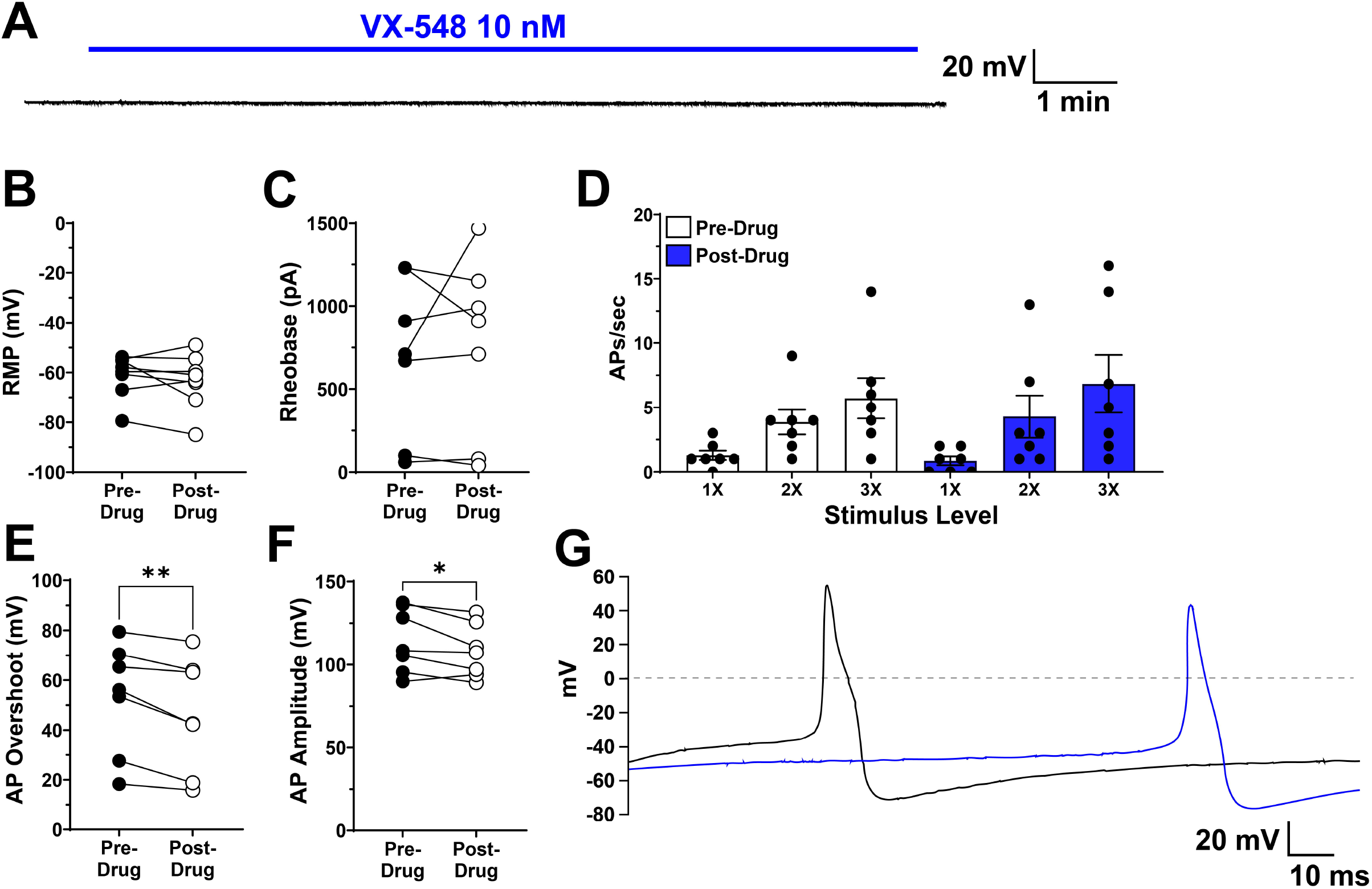
Suzetrigine alters action potential characteristics in MDA cancer patient DRG neurons without ectopic activity. In DRG neurons from MDA patients without spontaneous AP firing at baseline (*n* = 8), suzetrigine (VX-548, 10 nM) was applied for 10 min. (A) A representative raw trace example showing no effect on resting membrane potential (RMP) during or after VX-548 treatment, as shown by the blue bar. Overall, suzetrigine failed to alter (B) RMP, *t*(7) = 1.01, *P*=0.348, or (C) rheobase *t*(6) = 0.50, *P*=0.637, and no changes in response to 1 s current stimulation at 1X-3X rheobase (D) was observed, *F*(1,6) = 0.34, *P*=0.583. However, (E) AP overshoot *t*(6) = 4.15, *P*=0.006, and (F) AP amplitude, *t*(6) = 2.45, *P*=0.0499) were significantly lower post-drug, as shown by the representative raw trace example in (G), where the pre-drug AP waveform is shown in black and the post-drug AP waveform is shown in blue. \**P*<.05, ** *P*<.01.

Given the results described above, non-SA neurons from STA donors were challenged using a more prolonged suzetrigine application of up to 75 min (10 nM, *n* = 15). Now, responses to both ramp and step current stimulus protocols decreased significantly during drug perfusion compared to bath perfusion alone (Fig.3A,B). Neurons dissociated with protocol 1 showed an increase in rheobase (Fig.3C), a decrease in AP amplitude, more depolarized AP threshold, and AP overshoot was reduced (Fig.3D-G). Likewise, neurons dissociated using protocol 2 (see methods) showed a decrease in discharges evoked using both ramp and step current injections (Fig.4A-C), as well as depolarized membrane potential, decreased AP amplitude, and more depolarized AP threshold (Fig. 4D-E). No change in AP rise, fall, or AHP were observed (Supplementary Fig.3).

**Figure 3.**
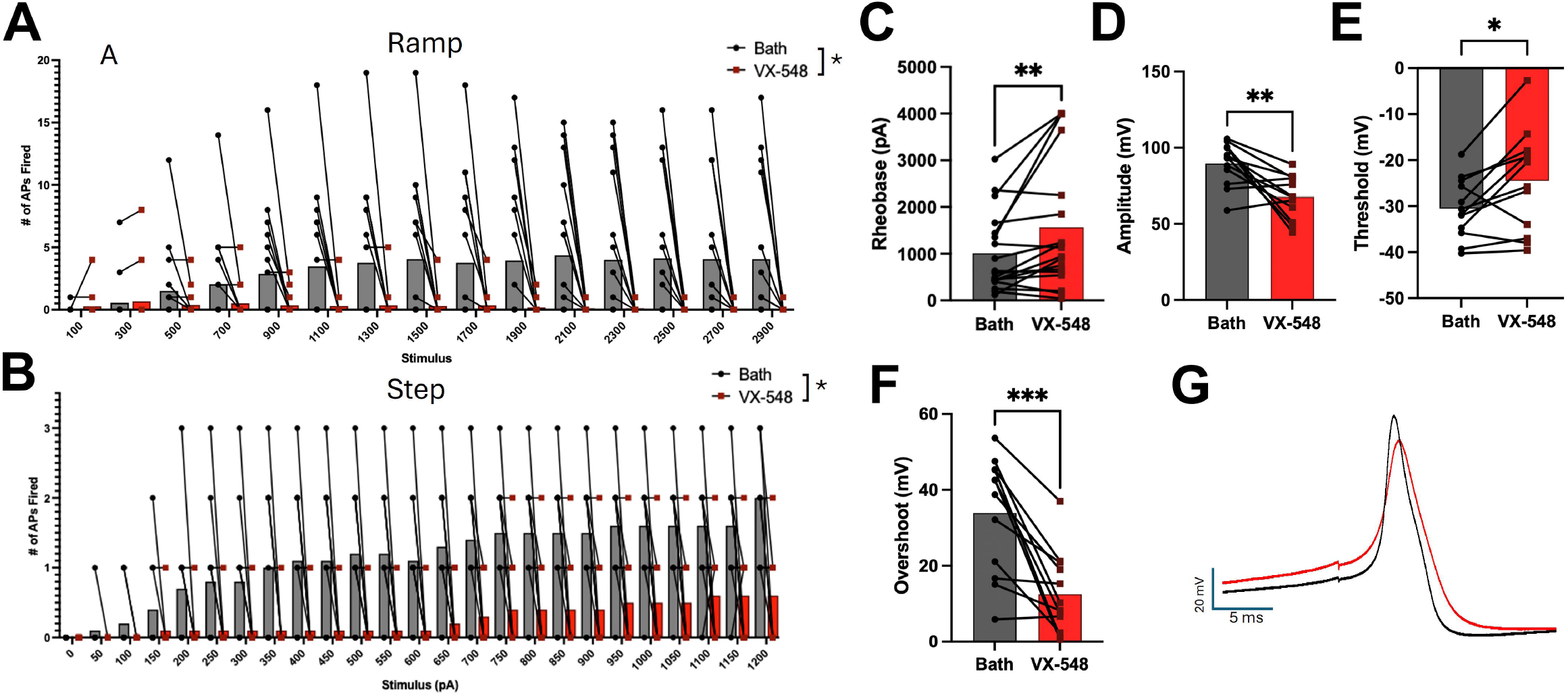
Suzetrigine rapidly reduces excitability in non-spontaneously active DRG neurons from STA donors (Protocol 1). (A) Evoked excitability was produced by a ramp stimulus applying a 1-100 pA ramp over 1 s, increasing the ramp end point by 200 pA each sweep. During VX-548 (10 nM) perfusion, the number of action potentials fired in each sweep significantly reduced relative to bath perfusion alone in the same cell (*n* = 15; *P* =0.0003, Wilcoxon matched pairs signed rank test). (B) VX-548 also decreased excitability to a step stimulus injecting 0 pA of current for 500 ms, increasing by 50 pA each sweep (*P* <0.0001, Wilcoxon matched pairs signed rank test). (B) In addition, VX-548 (10 nM) also increased the rheobase of hDRG neurons after less than 3 minutes of perfusion (*P* =0.0066). Among action potential characteristics, VX-548 decreased the (C) amplitude of the action potential at rheobase (*P* =0.0018), (D) shifted the action potential threshold to a more depolarized membrane voltage (*P* =0.0181), and (F) VX-548 decreased the overshoot of the action potential at rheobase (*P*=0.0003), as shown in (G) by the representative raw trace example of a cell with bath perfusion (black) and VX-548 perfusion (red).

**Figure 4.**
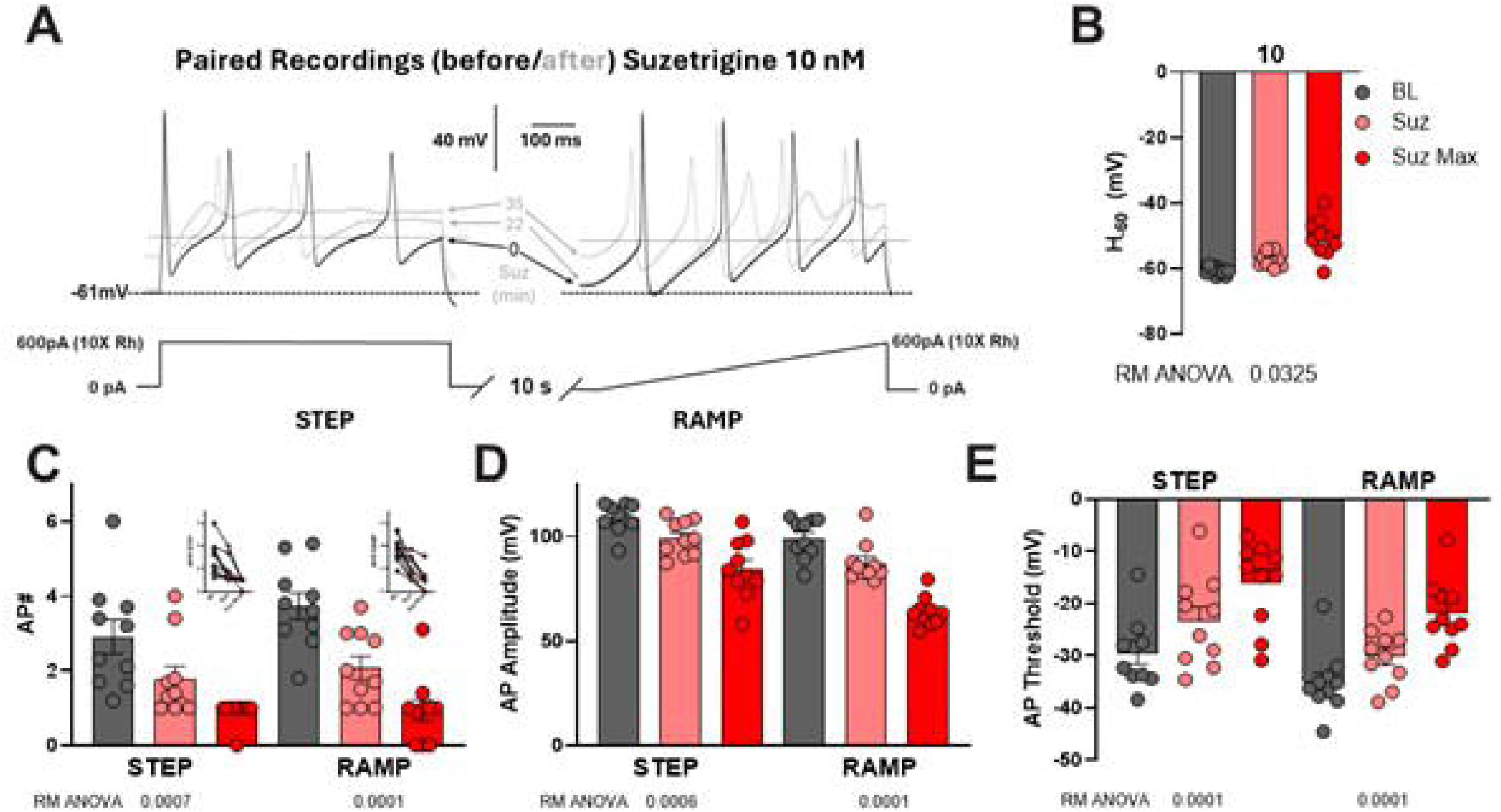
Suzetrigine decreased responses to step and ramp current injections. In DRG neurons from STA donors using dissociation Protocol 2 (*n* = 10), current injections above rheobase combining step and ramp protocols in the same sweep were applied before and after application of suzetrigine (VX-548, 10 nM). Drug was infused and testing protocols repeated until a change in response was observed (typically 5-30 min but up to 75 min), as shown in the representative raw trace in (A). The dashed line and the dotted line correspond to the baseline membrane potential and the threshold for the first AP, respectively. Below, step and ramp protocols separated by 10 s and the rheobase for this cell are shown. (B) Increased membrane potential from holding at −60 mV induced by suzetrigine. (C-D) Summary data for the effect of suzetrigine on APs. Number of APs fired before, at the midpoint of drug infusion, and at the maximum drug effect are presented in (C), showing significantly fewer APs firing after suzetrigine treatment. Suzetrigine also decreased action AP amplitude (D) and depolarized the firing threshold (E).

## Discussion

There are two main findings in this work. First, the FDA-approved Na_V_1.8 channel inhibitor suzetrigine readily suppressed on-going AP discharges in neurons from both patient and organ donor pain-associated DRG as well as in neurons from dermatomes without a pain history. Second, suzetrigine suppressed evoked activity in non-SA neurons but consistently only with more prolonged exposure than what suppressed on-going SA. A major strength of this work is that the results were found across DRG from different sources (patient and organ donors) and using multiple dissociation techniques, suggesting excellent generalizability of the findings regardless of these variables in source and preparation. There were some modest differences in effect of suzetrigine on SA neurons from patients versus donors in that suppression of SA in the patient DRG was associated with membrane hyperpolarization, decreased AP overshoot and AP width at 0 mV, but this was not as pronounced in the organ donor neurons. On-going activity was recoverable in some of the patient neurons after washout, but not seen in the organ donor neurons, which were recorded for a briefer period after the onset of drug effects. Given that SA in the patient donors have been clearly linked to the occurrence of on-going pain, the results here suggest that suzetrigine should likely prove efficacious in the management of this common feature of neuropathic pain^22,23^. Moreover, since the etiology of the pain-associated SA differed between the two sites involved, spinal nerve compression in the patient donors, versus diabetic neuropathy or unknown origin in the organ donor set, this also suggests that suzetrigine could prove effective on multiple types of on-going neuropathic pain.

Suzetrigine had little effect when used with a fixed application time on non-SA neurons from patients. Decreased AP overshoot was again observed, but there were minimal effects on RMP or current thresholds. However, when suzetrigine was tested using more prolonged application times-to effect of up to 75 min non-SA neurons from STA donors the drug caused increased current thresholds and decreased responses to sustained ramp and step current stimulation, AP overshoot and AP amplitude; as well as depolarized AP thresholds. These results are in agreement with a previous report showing an inhibition of evoked AP firing in non-SA DRG neurons from organ donors^8^. It is most likely that the less robust effects observed in non-SA neurons from MDA cancer patients were due to the differences in drug application time. However, it is also possible that differences in culture methodology, differential Na_V_1.8 expression based on pain history could also have contributed^15,16^. The DRGs from MDA were collected from patients who were older on average than the organ donors (63.2±1.7 vs 36.4±4.9 yo) and with highly variable cancer and treatment histories. Further, all MDA patient DRGs were from thoracic dermatomes, while STA donor DRGs were from lumbar levels. Different dissociation protocols could also result in different subpopulations of neurons viable for recording or shifts in the proportion of Na_V_1.8-expressing neurons over time^25^. MDA experiments were performed 12-36h after plating, while UTD recording took place across a much longer *in vitro* period (3-28 days). Further study is needed to determine how these variables impact Na_V_1.8 expression and function in cultured DRG neurons.

The impacts of suzetrigine on AP overshoot and amplitude were expected due to Na_V_1.8’s role as an overshoot channel in AP electrogenesis^26^, while the AP width and firing threshold may have been altered due to hyperpolarization of the membrane potential (reaching less than −90 mV in some neurons) affecting the function of voltage-gated potassium and calcium channels or hyperpolarization-activated cyclic nucleotide calcium (HCN) channels^27–29^. No change in the number of APs fired in response to 1-3x current stimulation were found for SA or non-SA neurons. Thus, it appears that the effects of this drug emerge primarily in active neurons in the MDA patient population. This is similar to a different Na_V_1.8 inhibitor, A-308467, which blocked AP firing in depolarized spontaneously active neurons and inhibited AP firing at threshold only when cells were depolarized via current injection to −40 mV, with minimal impacts at −53 mV^30^.

Overall, our studies suggest a role for Na_V_1.8 inhibition in silencing ectopic discharge, including that induced by spinal cord/root compression, and attenuating hyperexcitability. In patients with neuropathic pain, this suppression of ectopic firing may correspond to relief of spontaneous pain, but there may not necessarily be changes to evoked pain, as suppression of evoked responses occurred only when sustained current stimulation was applied, suggesting that Na_V_1.8 may play a role in repetitive firing, but not the initiation of AP firing. This has previously been reported in clinical trials, where suzetrigine pain scores were lower than placebo, but pain was not eliminated^9^. For a potential analgesic drug, activity-dependent effects are ideal, as non-sensitized neurons can continue to function normally while ectopic activity in sensitized neurons is suppressed. Treatment approaches that incorporate more targeted treatments such as suzetrigine for pain could reduce reliance on opiates with more severe side effect profiles and improve quality of life in patients.

## Acknowledgments

The authors would like to thank the patients, donors, and their families for their participation, members of the Southwest Transplant Alliance and Price and Dussor labs at UTD for supporting the tissue recovery work.

## Funding

This research was supported by NIH 5R01NS111929 (PMD, TJP) and NIH 5U19NS130608 (TJP, PMD, MC).

## Competing interests

The authors report no competing interests.

## Supplementary material

Supplementary material is available at *Brain* online.

## Notes

### Competing Interest Statement

The authors have declared no competing interest.

