## Supplemental Tables, Figures and Methods for "Suzetrigine (VX-548) exhibits activity-dependent effects on human dorsal root ganglion neurons"

**Supplementary Material**

**Supplementary Methods and Materials**

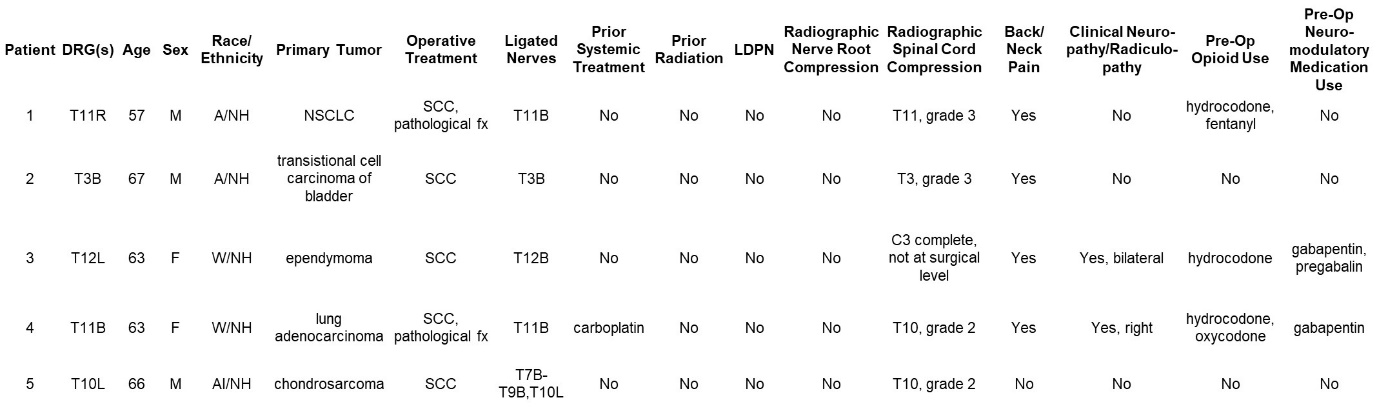

**Supplementary Table 1. Demographics and detailed clinical data for the MDA patient cohort.**

**Sex: M = male, F = female**

**Race/Ethnicity: A = Asian, AI = American Indian, W = White, H = Hispanic/Latino, NH = Non-Hispanic/Latino**

**Primary tumor: NSCLC = non-small cell lung cancer**

**Operative Indication: SCC = spinal cord compression, fx = fracture**

**LDPN = length dependent peripheral neuropathy**

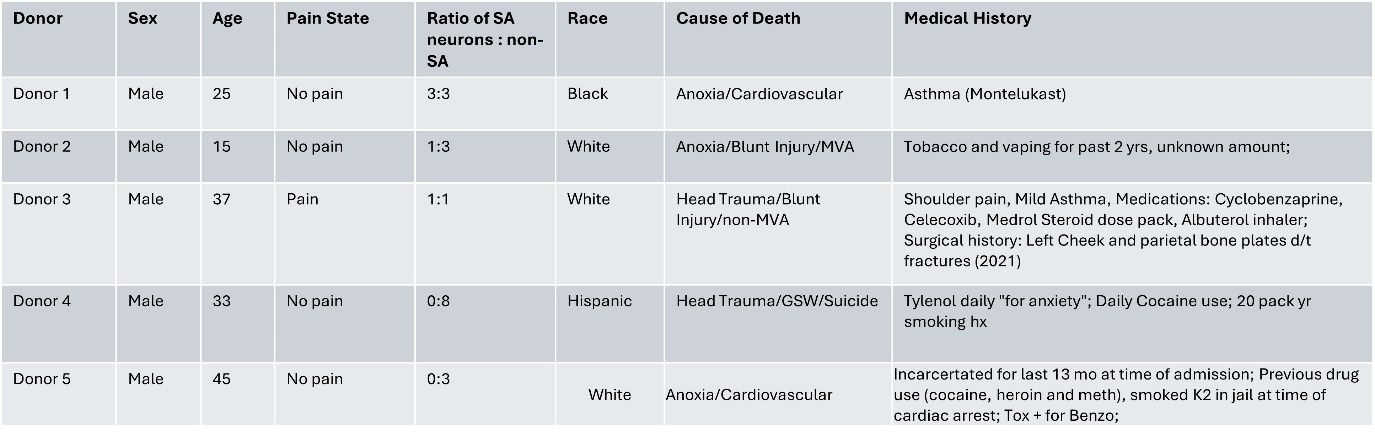

**Supplementary Table 2. Demographics and detailed clinical data for STA donors (Protocol 1)**

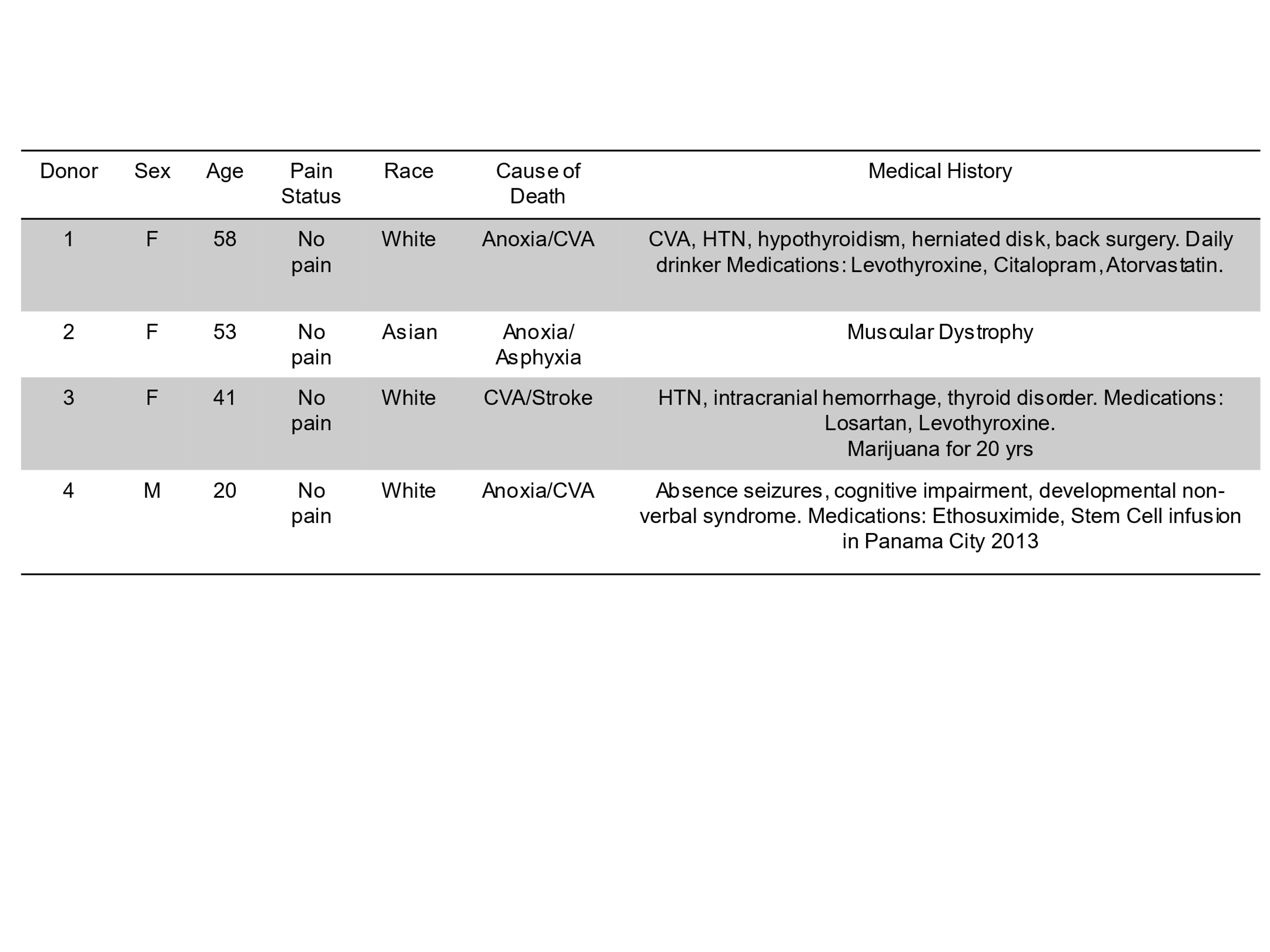
**Supplementary Table 3. Demographics and detailed clinical data for STA donors (Protocol 2)**

**Whole-cell recording of dissociated human DRG neurons**

***MDA Thoracic Vertebrectomy Patients***

Briefly, dissociated DRG neurons were transferred to a recording chamber perfused with oxygenated (95% O_2_ + 5% CO_2_) extracellular solution containing 117 mM NaCl, 3.6 mM KCl, 1.2 mM NaH_2_PO_4_·H_2_O, 2.5 mM CaCl_2_, 1.2 mM MgCl_2_, 25 mM NaHCO_3_ and 11 mM glucose adjusted to pH 7.4 with NaOH. Glass-micropipettes (8-10 MΩ) were filled with an internal solution with 125 mM KCl, 15 mM K-gluconate, 5 mM Mg-ATP, 0.5 mM Na2GTP, 5 mM HEPES, 2 mM MgCl2, 5 mM EGTA, and 0.5 mM CaCl_2_ adjusted to pH 7.4 with KOH. Whole-cell patch clamp electrophysiology was performed using a Multiclamp 700B amplifier (Molecular Devices), Digidata 1440 digitizer (Molecular Devices) and pClamp 10 acquisition software (Molecular Devices). Only DRG neurons with a resting membrane potential of at least -40 mV, stable baseline recordings, and evoked spikes that overshot 0 mV were used for further experiments and analysis. Series resistance (Rs) was compensated to above 70%. All recordings were made at room temperature. In voltage clamp, neurons were held at −70 mV, and activation was evoked with a 15 ms step to potentials ranging from −60 to +80 mV in 10 mV increments to ensure cell health. The DRG neurons were then held at 0 pA in current clamp to determine the presence or absence of spontaneous action potential for 5 min. DRG neurons were then held at 0 pA, and action potentials were evoked using a series of 500 ms depolarizing current injections in 10 pA steps starting from -50 pA. The current that induced the first action potential was defined as the current threshold (1X rheobase). Neurons were then stimulated with 1 s current injections at 1×, 2×, and 3× rheobase. VX-548 diluted to a 10 nM working concentration in extracellular solution was perfused into the recording chamber with a flow rate of 2 mL/min. Drug was applied until APs stopped firing (3-20 min) in neurons with ectopic discharge, followed by a washout period (2-11 min) before re-testing for rheobase and responses to current injections at 1×, 2×, and 3× rheobase. In non-SA neurons, drug was applied for 10 min, followed by a 5 min washout period before re-testing rheobase and responses to current injections at 1×, 2×, and 3× rheobase. In some neurons, capsaicin was used to assess whether neurons were TRPV1 positive. Capsaicin (Sigma) stock solution was prepared in ethanol and diluted in extracellular solution to a working concentration of 100 nM and was applied for 1-5 min followed by a washout period. Two non-SA neurons were challenged with capsaicin prior to VX-548 treatment, did not respond, and so a 30 min washout was implemented and baseline testing repeated prior to VX-548 treatment. Data analysis was conducted using Clampfit 11.1 (Molecular Devices, San Jose, CA) software.

Changes in AP firing frequency during and after Suzetrigine treatment were analyzed using repeated measures ANOVA with time bin as the within-subjects variable and Bonferroni post-hoc tests. The number of APs in five one-minute time bins (last minute of baseline, first and last minute of drug infusion, first and last minute of washout) were used for this analysis. Changes in RMP, rheobase, and AP characteristics pre- and post-drug were analyzed using dependent t-tests. Responses to current stimulation at 1X, 2X, and 3X rheobase were expressed as the number of AP fired per second and analyzed using repeated measures ANOVA and Bonferroni post-hoc tests to compare pre- and post-drug responses.

***UTD Donors (Protocol 1)***

Whole-cell patch clamp electrophysiology was conducted using a Multiclamp 700B (Molecular Devices, San Jose, CA) amplifier, a Digidata 1550 digitizer (Molecular Devices, San Jose, CA), and pClamp10 acquisition software (Molecular Devices, San Jose, CA). After day 3 in vitro, hDRG neurons plated on glass coverslips were transferred from media to bath solution (135 mM NaCl, 10 mM Glucose, 10 mM HEPES, 5 mM KCl, 2 mM CaCl_2_, 1 mM MgCl_2_, 20 mM sucrose, with a pH of 7.4 and osmolarity ranging from 310-320) and all data was collected within 90 minutes of transfer. Glass micropipettes (outer diameter 1.5mm: inner diameter, 0.89 mm; BF150-110-10, Sutter Instruments) were pulled using a PC-100 puller (Narishige, Amityville, NY) and fire polished to the resistance of 1-3 MΩ using a microforge (MF-83, Narishige). The pipettes were then filled with pipette solution (135 mM KCl, 10 mM HEPES, 4 mM ATP-Mg, 0.9 mM GTP-Na, 0.5 mM EGTA, 5 mM NaCl with a pH of 7.4 and osmolarity ranging from 305-320) and pipette resistance was offset. The pipette was then attached to the cell and suction applied until a giga-ohm seal was achieved. The membrane was then ruptured, fast and slow capacitance were offset, and whole cell mode was achieved. A -70 mV to 0 mV to -70 mV voltage step was applied to ensure cell health. Bath perfusion was initiated and the amplifier was switched to current clamp mode. In current clamp mode, a 90 s recording with no stimulus is taken to assess spontaneous activity prior to a current step injecting 0 pA of current for 20 ms with a delta of 10 pA was applied until the cell fired the first action potential to measure rheobase. Next, a ramp current injection was applied to the cell ramping from 0-100 pA over 1 s, with a delta of 200 until 2900 pA. Finally a step current injection of 0 pA over 500 mS with a delta of 50 pA was injected to 2100 pA. Perfusion was then switched to bath containing 10 nM VX-548 and all recordings are taken again. Data analysis was conducted manually using Clampfit 11.4 (Molecular Devices, San Jose, CA) software. Data are expressed as mean ± SEM.

***UTD STA Donors (Protocol 2)***

Recordings were done 3 or more days after plating. The external solution contained (in mM): 125 NaCl, 3.6 KCl, 2 NaH_2_PO_4_, 2.5 CaCl_2_, 2 MgCl_2_, 25 NaHCO_3_, and 11 dextrose. Whole-cell patch clamp was performed using borosilicate capillaries pulled with a P-97 flaming-brown micropipette puller (Sutter Instruments). The pipettes had a resistance of 1.5-4 MΩ, when using an internal solution containing (in mM): 120 K-Glutamate, 2 KCl, 8 NaCl, 0.2 EGTA, 14 Na2-Phoshphocreatine, 2 Mg-ATP, 0.3 Na-GTP and 10 HEPES-K (pH 7.3, 295 mOsm). Recordings were obtained using an Axopatch 200B amplifier (Molecular Devices). Neurons were visualized using a Nikon Eclipse Ti inverted microscope equipped with Nikon Advanced Modulation Contrast. Acquisition was done at 10-20 kHz and data was filtered at 5 kHz and analyzed offline using Clampfit Analysis Suite 11 (Molecular Devices), GraphPad Prism 11, and Microsoft Excel 2015. In whole-cell configuration, cells were held in voltage clamp mode at near resting potential for at least 5 min to allow for dialysis of the pipette internal solution. Afterwards, when needed, cells were held at -60 mV (H_-60_) by injecting current of the appropriate sign and amplitude. For rheobase evaluation, first, a gross determination of action potential firing was made using 10-200 pA incremental steps (or ramps; 500 ms duration), and then a baseline subthreshold current with incremental steps (or ramps) of 1/10th of the apparent Rh in successive sweeps were used. The rheobase was considered the step (or ramp) at where the first action potential (AP) was fired and where APs were consistently fired in the following steps. The acute effect of suzetrigine on evoked firing was evaluated using both step and ramp protocols separated by 10 seconds within the same sweep and repeated at 0.033 Hz and where the stimulation intensity was increased within the range of 2X to 10X Rh to achieve multi-firing. Ten baseline sweeps were recorded before 10-20 nM suzetrigine was added to the batch solution. Recordings continued until an effect was seen or up to 75 min, whatever was first (usually between 10-30 min).

**Supplementary Results**

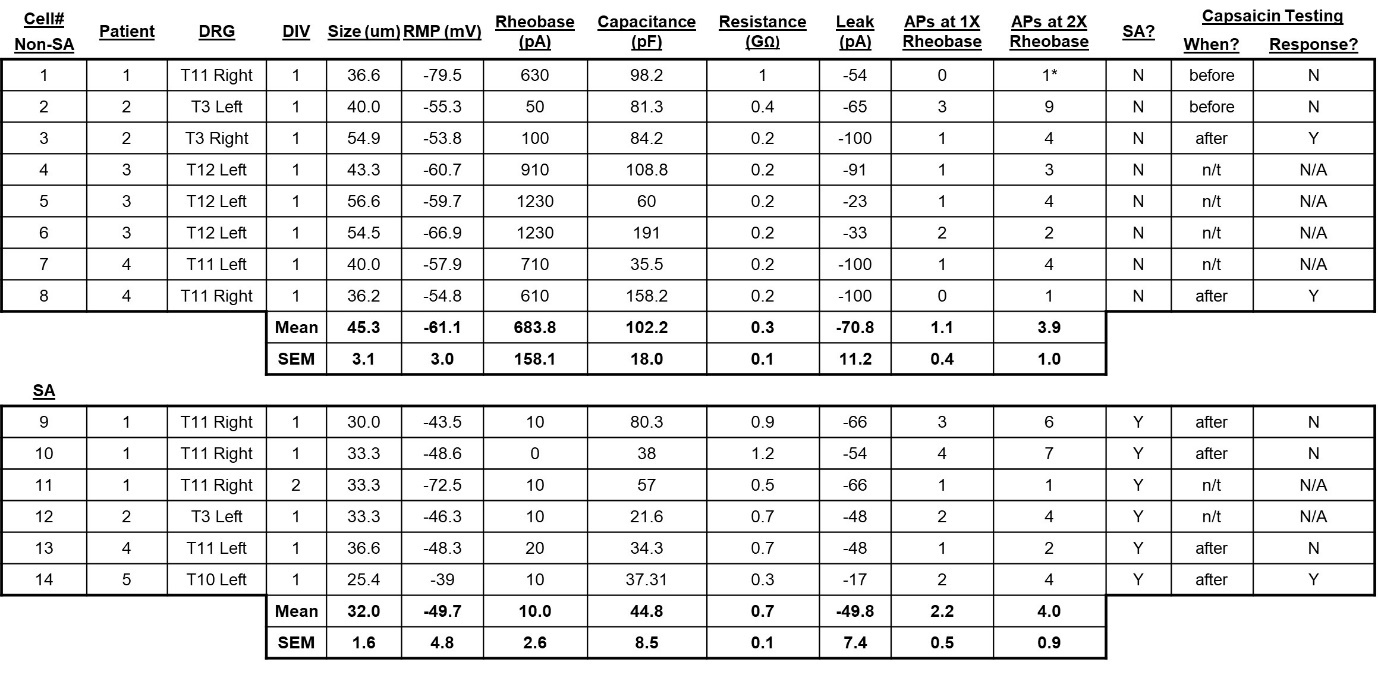

**Supplementary Table 4.** *MDA patient DRG neuron characteristics*. Details for DRG neurons without spontaneous activity at baseline (non-SA) appear at the top of the table and neurons with spontaneous activity (SA) are shown at the bottom. Neurons were challenged with capsaicin (100 nM) for up to 3 min before testing VX-548, after testing VX-548, or not tested (n/t). Y = positive response to capsaicin, N = no response to capsaicin, N/A = not applicable. * = rapidly accommodating neuron

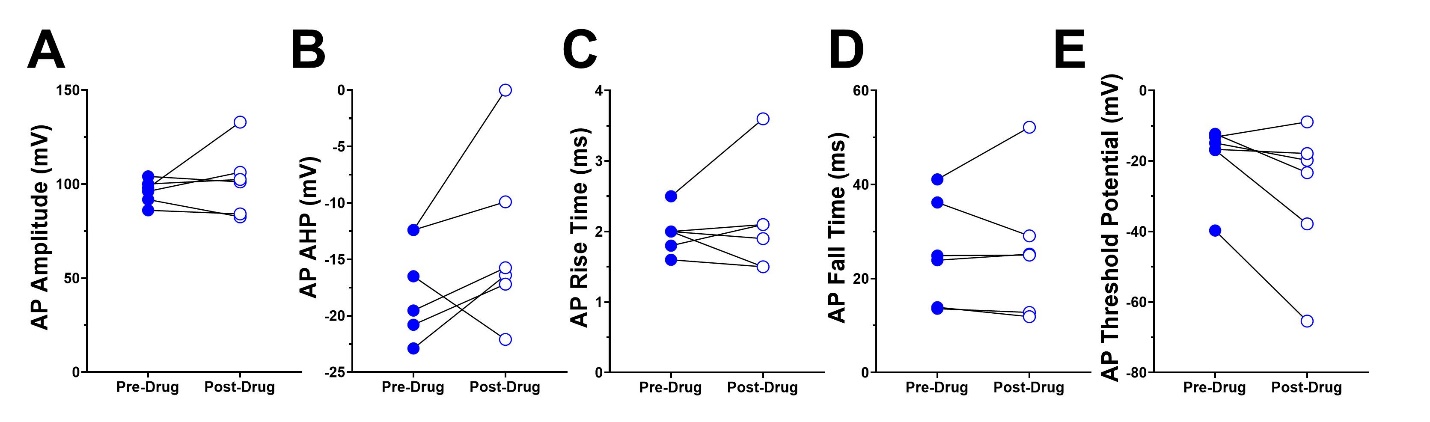

**Supplementary Figure 1.** In spontaneously active DRG neurons from MDA cancer patients, action potential (AP) characteristics not changed by treatment with 10 nM VX-548 included (A) AP amplitude (t_5_ = 0.87, p=0.426), (B) afterhyperpolarization (AHP; t_5_ = 1.62, p=0.167), (C) rise time (t_5_ = 0.60, p=0.574), (D) fall time (t_5_ = 0.18, p=0.867), and (E) threshold potential (t_5_ = 2.09, p=0.091).

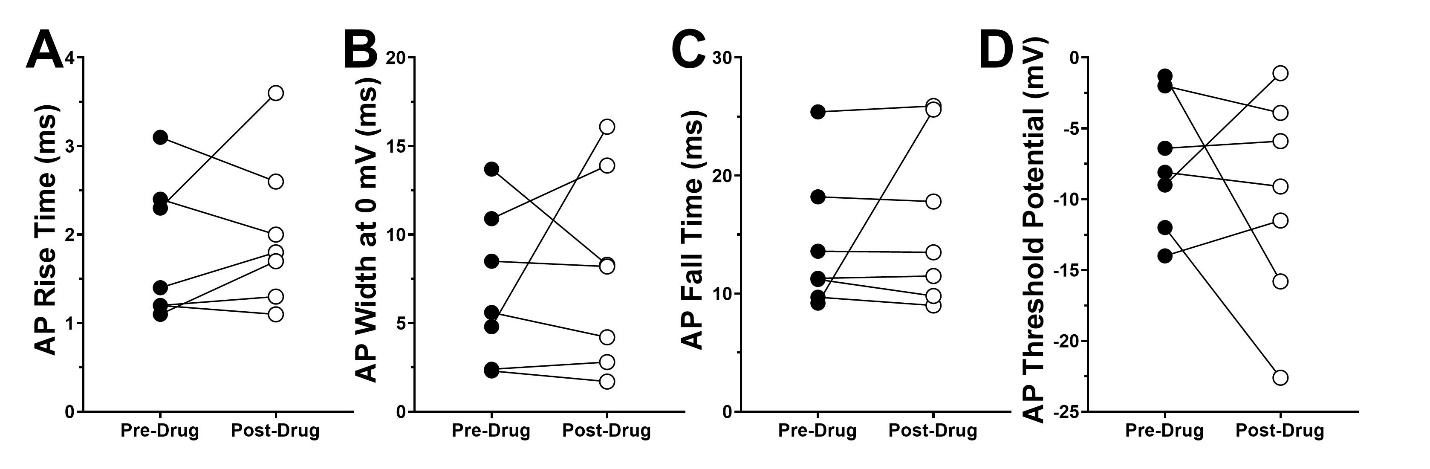

**Supplementary Figure 2**. In DRG neurons from MDA cancer patients without ectopic activity, action potential (AP) characteristics not changed after 10 min treatment with 10 nM VX-548 include (A) AP rise time (t_6_ = 0.84, p=0.431), (B) AP width at 0 mV (t_6_ = 0.51, p=0.628), (C) AP fall time (t_6_ = 0.86, p=0.421), and (D) AP threshold potential (t_6_ = 0.84, p=0.433). Summary data for AP afterhyperpolarization (AHP) is not shown as only 2 of 8 neurons demonstrated AP AHP at baseline.

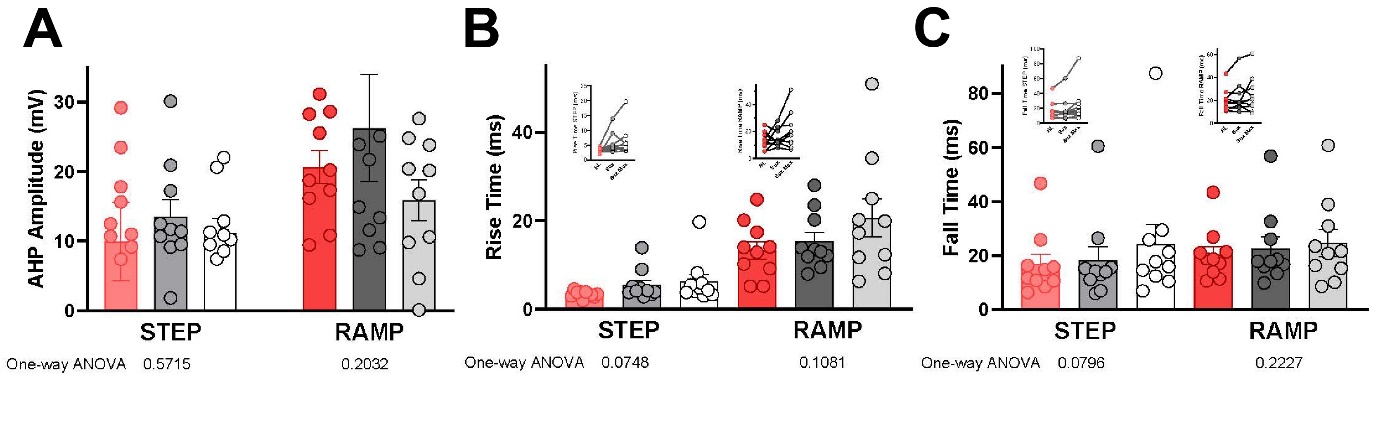

**Supplementary Figure 3**. In DRG neurons from STA donors (Protocol 2), treatment with 10 nM VX-548 did not alter action potential (AP) rise time (A), fall time (B), or afterhyperpolarization (AHP, C).
